# Brain structure mediates the association between height and cognitive ability

**DOI:** 10.1101/183525

**Authors:** Eero Vuoksimaa, Matthew S. Panizzon, Carol E. Franz, Christine Fennema-Notestine, Donald J. Hagler, Michael J. Lyons, Anders M. Dale, William S. Kremen

## Abstract

Height and general cognitive ability (GCA) are positively associated, but the underlying mechanisms of this relationship are unclear. We used a sample of 515 middle-aged male twins with structural magnetic resonance imaging data to study if the association between height and cognitive ability is mediated by cortical size. We used genetically, ontogenetically and phylogenetically distinct cortical metrics of cortical surface area (SA) and cortical thickness (CT). Our results indicate that the well-replicated height-GCA association is accounted for by individual differences in total cortical SA (highly heritable metric related to global brain size), and not mean CT, and that the genetic association between SA and GCA underlies the phenotypic height-GCA relationship.

## Introduction

Both height and general cognitive ability (GCA) are predictors of health outcomes, and there is a positive height-GCA correlation of .10 -.20 (Keller et al., 2013; Marioni et al., 2014; Silventoinen et al., 2012). There has been widespread interest in the association between height and cognitive ability from the development of preterm infants (Sammallahti et al., 2014) to aging-related dementia (Russ et al., 2014). Height is highly heritable but is also considered to reflect early life events and is often regarded as a proxy for early brain development. Still, the neural substrate of the height-cognitive ability association is unclear. Understanding its neural underpinnings may shed light on pathways to normative and abnormal development and aging.

Both height and GCA have substantial heritability and their association is largely due to shared genetic effects (Marioni et al., 2014; Silventoinen et al., 2012). Both are positively correlated with brain volume (which is also highly heritable), and these associations are also largely due to shared genetic effects (Posthuma et al., 2000; Posthuma et al., 2002). Studies have examined head size or global brain measures with respect to height (Adams et al., 2016; Taki et al., 2012). Association between brain volume and GCA remains when controlling for height (Andreasen et al., 1993) and a map-based study in children using cortical volume (CV) measures of gray matter indicated some overlapping regional brain volume-height and brain volume-GCA correlations (Taki et al., 2012). Still, no studies have investigated if cortical structure mediates the association between height and GCA.

During the past decade, studies of GCA-cortical relationships have moved from CV measures to investigations of cortical thickness (CT) and surface area (SA) separately (Panizzon et al., 2009; Vuoksimaa et al., 2015), two metrics that are phenotypically and genetically uncorrelated at the global level (Panizzon et al., 2009; Vuoksimaa et al., 2015). However, we are unaware of any investigations of the role that CT and SA play in the relationship between height and GCA. A twin design could further explicate the genetic/environmental underpinnings of these associations.

We examined 515 middle-aged male twins (51–60 years) with data on GCA, height and neuroimaging with 1.5T MRI to investigate whether cortical structure mediates the height-cognitive ability association and the genetic underpinnings of those relationships. We hypothesized that the height-GCA association would be accounted for by cortical SA, rather than CT, because SA is much more strongly related to cortical, whole brain, and intracranial volumes, and GCA in comparison to CT (Vuoksimaa et al., 2015).

## Methods

### Participants

Participants were 534 middle-aged men (51 – 60 year olds; M = 55.7; SD = 2.6) from the Vietnam Era Twin Study of Aging (VETSA) (Kremen et al., 2006; Kremen et al., 2013). After quality control, we had measures of total cortical grey matter surface area and cortical thickness on 515 participants (Vuoksimaa et al., 2015). The sample consisted of 131 monozygotic (MZ) and 96 dizygotic (DZ) full twin pairs and 61 individual without a co-twin. Our sample of over 500 participants is well powered to detect small height-GCA correlation of about .10 -.20 as suggested by earlier literature.

Zygosity determination was based on 25 microsatellite markers for majority of the participants (92%) and on questionnaire and blood group for the remaining participants. Questionnaire and blood group based classification had 95% agreement with the DNA-based method.

Data on general cognitive ability (GCA) and neuroimaging were collected on back-to-back days. Height was assessed in stocking feet with a stadiometer, rounded to the nearest half inch and then converted to cm. Height and GCA data were acquired on the same day. Data were collected at two sites: University of California, San Diego and Boston University (neuroimaging at the Massachusetts General Hospital). Ethical approval was obtained from all participating institutions and all participants gave written informed consent.

### General Cognitive Ability (GCA) Measure

GCA was measured with Armed Forces Qualification Test (AFQT). The AFQT is a 50-min paper-and-pencil test consisting of items measuring verbal ability (vocabulary); arithmetic, spatial processing (mental folding and unfolding of boxes); and reasoning about tools and mechanical relations. The total score on the AFQT has a correlation ∼0.85 with Wechsler IQ and in the VETSA sample test–retest reliability was 0.74 over 35 years and 0.73 over 42 years (Lyons et al., 2009; Lyons et al., 2017). AFQT yields a percentile score but in the analyses we used a variable whereby the percentile scores were transformed into their normal deviates (Lyons et al., 2009).

### Image Acquisition and Processing

Detailed descriptions of MRI image acquisition and processing can be found in Kremen et al.(Kremen et al., 2010) and Eyler et al.(Eyler et al., 2012). Briefly, images were acquired on Siemens 1.5 T scanners. Two sagittal T1-weighted MPRAGE sequences were employed with a TI = 1000 ms, TE = 3.31 ms, TR = 2730 ms, flip angle = 7°, slice thickness = 1.33 mm, and voxel size 1.3 × 1.0 × 1.3 mm. To increase the signal-to-noise ratio, the two MPRAGE acquisitions were rigid-body registered to each other (motion corrected) and then averaged.

Volume segmentation (Fischl et al., 2002; Fischl et al., 2004a) and cortical surface reconstruction (Dale and Sereno, 1993; Dale et al., 1999; Fischl et al., 1999; Fischl et al., 2004b) were performed using the publicly available FreeSurfer software package. The three-dimensional cortical surface was reconstructed to measure thickness and area at each surface location or vertex using a semi-automated approach in the FreeSurfer software package. The cortical surface was covered with a triangular tessellation, which was then smoothed to reduce metric distortions. A refinement procedure was then applied to obtain a more accurate representation of the gray/white boundary. This surface was then deformed outwards to obtain an explicit representation of the pial surface. The resulting cortical surface model was manually reviewed and edited for technical accuracy. Minimal manual editing was performed according to standard, objective editing rules. These semi-automated measures have a high correlation with manual measures in vivo and ex vivo (Fischl and Dale, 2000; Walhovd et al., 2005).

Total surface area was calculated as the sum of the areas of all vertices across the cortex. Cortical thickness was calculated as the average distance between the gray/white boundary and the pial surface within each vertex (Fischl and Dale, 2000). Mean cortical thickness was calculated as the average thickness of the whole cortex, weighted by surface area of each vertex.

### Bootstrapped mediation models

We ran a multilevel mediation model in Stata (ml_mediation) with family as a cluster variable (i.e., robust standard errors accounting for the clustered within-family twin data). GCA was a dependent variable, total SA was a mediator variable and height was independent variable. The effects of age and scanner (different scanners were used in Boston and San Diego data collection sites) were regressed out of total SA before the analysis.

All variables were standardized with a mean of 0 and standard deviation of 1. A bootstrapped method with 1000 replications was used to derive 95% confidence intervals. Path estimates that do not include zero in the confidence intervals were considered as statistically significant. We ran additional mediation model whereby height and total SA variables were considered as mediator and independent variables, respectively.

### Biometrical genetically informative twin analyses

By studying monozygotic (MZ) and dizygotic (DZ) twins it is possible to estimate the relative proportion of genetic and environmental influences on a given phenotype. In a classical twin design, the variance of a phenotype is decomposed into additive genetic influences (A), common environmental influences (C), and unique environmental influences (E). This model is referred as an ACE model. A, C, and E are latent variables and they do not refer to specific genes or specific environmental events. Genetic effects from these models can be interpreted as a heritability estimate which tells how much of the variation between individuals in a given trait in a given population is due to genetic effects.

The A effects correlate 1.0 in MZ twin pairs, who are assumed to share all of their genes, whereas these effects correlate 0.5 in DZ twin pairs who share on average half of their segregating genes. C effects correlate 1.0 both in MZ and DZ twin pairs and refer to all environmental effects that make co-twins similar to each other. The E effects refer to all environmental effects that make co-twins different from each other and are therefore uncorrelated both in MZ and DZ twin pairs. The E component also includes measurement error. Analyses were performed using the maximum-likelihood based, structural equation modeling software Mx (Neale et al., 2004).

To determine the relative contribution of genetic and environmental influences on both the individual measures and the covariance between measures, we fit Cholesky decompositions to the data. All variables were normally distributed, meeting a basic assumption of these parametric analyses. The effects of age and scanner were regressed out of the cortical measures prior to all analyses.

The univariate ACE model is easily extended to a trivariate scenario in which the sources of genetic and environmental covariance can also be examined. We began with trivariate models that included A, C and E variance components for GCA, total SA, and height. This starting point was called as the “ACE–ACE–ACE” Cholesky. Next, a reduced ACE-AE-AE Cholesky decomposition in which the C effects for total SA and height were fixed to be zero was tested relative to the full ACE-ACE-ACE Cholesky decomposition. We have previously used the Cholesky decompostions in the context of GCA, total SA and mean CT. Our previous work (Vuoksimaa et al., 2015) showed that C effects were 0.17 for GCA, but the lower bound of the 95% confidence interval included zero indicating that C effects could also be fixed to zero in the case of GCA. However, we kept the C component for GCA in the subsequent models because we believe that the magnitude of C effect was too large to be fixed at zero. It is important to note that all covariance between twins is accounted either by A or C effects. Thus, fixing C to zero would inflate the magnitude A. Our approach of keeping the C effect for GCA is therefore a conservative approach which reduces the possibility that genetic covariation between traits is confounded by the C effects.

We computed phenotypic, genetic and unique environmental correlations between GCA, total SA and height. Phenotypic correlations are the standard correlations; they represent a composite of the shared genetic and environmental sources of variance (Neale and Cardon, 1992). Genetic correlations represent only the shared genetic variance between variables, and unique environmental correlational represent the shared individual-specific environmental variance between variables (Neale and Cardon, 1992).

Model comparisons were based on the likelihood-ratio χ2-test, which is calculated as the change in -2 log likelihood (–2LL) from the ACE–ACE-ACE to the ACE-AE-AE Cholesky or from the ACE-AE-AE to the ACE-AE-AE Cholesky whereby the genetic correlation between two variables is fixed to be zero, and is distributed as a χ2 with degrees of freedom equal to the difference in parameters between the models. Nonsignificant p-values (>0.05) indicate that the reduced solution does not yield a significant change in the model fit and therefore provides an essentially equally good fit to the data while using fewer parameters.

## Results

Measured height (M=177cm, SD=6.7cm) was positively correlated with GCA (r=.10, p=.022), CV (r=.24, p<.0001) and SA (r=.25, p<.0001), but not with CT (r=.04, p=.41). We previously reported the sample description and descriptive statistics for SA, CT and GCA in (Vuoksimaa et al., 2015) where we also showed that the association between CV and GCA is driven by SA rather than. Since both height and GCA were associated with SA but not with CT, we used only SA in subsequent analyses.

Height was significantly associated with GCA (Fig. 1A). Cortical SA significantly mediated the association between height and GCA (Fig. 1B). In the mediator model, the direct effect of height on GCA (β =.080, 95%CI= -.044; .205) was not significant (Fig. 1, Supplementary Table 1). The indirect effect of height on GCA through total SA (β =.048, 95%CI= .014; .082) accounted for 37% of the total effect of height on GCA (Fig. 1B).

**Fig.1.**
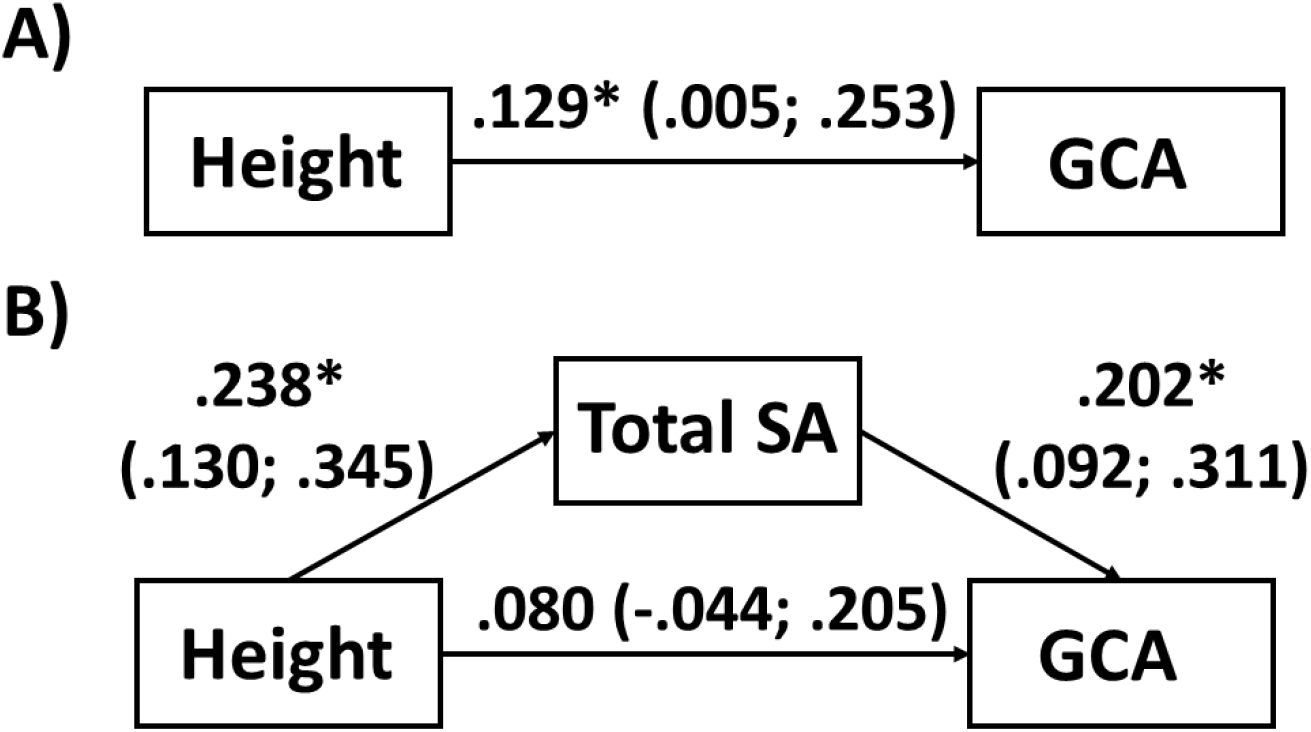
Associations between height, total surface area (SA) and general cognitive ability (GCA). Path estimates with 95% confidence intervals in parentheses. A) Path estimates for the effect of height on GCA. B) Mediation model path estimates of direct and indirect effects from height to GCA. *=statistically significant path estimates. All variables standardized (M=0, SD=1).

**Table 1.** Trivariate model fitting results for general cognitive ability (GCA), total cortical surface area (SA), and height. Note. Model 1 is the comparison model for Model 2; Model 2 is the comparison model for Models 3 and 4. A=additive genetic variance; C=common environmental variance; E=unique environmental variance; rg=genetic correlation; -2ll=-2 log likelihood; LRT=likelihood-ratio χ2 –test; df=degrees of freedom; Δdf=change in degrees of freedom. Model 4. includes A, C & E effects for GCA, A & E effect for SA, A & E effects for height and significant SA-GCA rg, significant SA-height rg, but no significant GCA-height rg. Best-fitting model shown in **bold** font.

| Model | -2ll | LRT | df | Δdf | p-value |
| --- | --- | --- | --- | --- | --- |
| 1. ACE-ACE-ACE Cholesky | 3569.826 |  | 1524 |  |  |
| 2. ACE-AE-AE | 3571.036 | 1.210 | 1529 | 5 | 0.944 |
| 3. ACE-AE-AE,<br>GCA-SA $r_g = 0$ | 3584.029 | 12.993 | 1530 | 1 | 0.001 |
| 4. **ACE-AE-AE,**<br>**GCA-Height $r_g = 0$** | **3574.666** | **3.631** | **1530** | **1** | **0.057** |

Note. Model 1 is the comparison model for Model 2; Model 2 is the comparison model for Models 3 and 4. A=additive genetic variance; C=common environmental variance; E=unique environmental variance; $r_g$ =genetic correlation; -2ll=-2 log likelihood; LRT=likelihood-ratio $\chi^2$ -test; df=degrees of freedom; Δdf=change in degrees of freedom. Model 4. includes A, C & E effects for GCA, A & E effect for SA, A & E effects for height and significant SA-GCA $r_g$ , significant SA-height $r_g$ , but no significant GCA-height $r_g$ . Best-fitting model shown in **bold** font.

We also ran a mediator model with SA as an independent variable and height as a mediator. In this model, the direct effect from SA to GCA was significant (β =.202, 95%CI= .092; .311) but the indirect effect of SA to GCA through height was nonsignificant (β =.019, 95%CIs= – .014; .052) (Supplementary Fig. 1).

Fixing C effects to zero for SA and height did not result in a poorer fit relative to a full ACE-ACE-ACE Cholesky where all variables included A, C and E effects (model 2 vs model 1in Table 1). Heritability estimates (95% CIs in parentheses) from the biometric trivariate Cholesky decomposition were: .59 (.33;.82) for GCA, .94 (.92;.96) for SA, and .93 (.91;.95) for height (from model 2 in Table 1). Constraining the GCA-SA genetic correlation to be zero did result in a poorer model fit (Table 1, model 3), but constraining the genetic correlation between height and GCA to be zero did not result in a poorer model fit (Table 1, model 4). In the best-fitting Cholesky decomposition (Table 1, model 4), phenotypic and genetic correlations between GCA and SA were .19 (.09;.28) and .22 (.09;.36), respectively.

## Discussion

The underlying mechanism of the well-replicated association between height and GCA has heretofore remained unclear. Our mediation models indicated that the height-GCA association is accounted for by brain size as characterized by individual differences in total cortical SA. Across adulthood, reductions in CV are related more to reductions in CT than SA, suggesting that SA may be a more stable cortical metric after childhood (Storsve et al., 2014). Although we did not have neuroimaging data in young adulthood and our finding of cortical SA mediating the effect of height on GCA was based on middle-age sample, other studies have indicated stable SA-GCA (Walhovd et al., 2016) and height-GCA (Harris et al., 2016) associations throughout life.

Both height-GCA and SA-GCA associations likely have origins in early development. Head growth in the first years of life reflects rapid growth in brain size, especially with respect to cortex. SA undergoes remarkable growth from 0 to 2 years with an average 1.8-fold expansion in the first year of life (Li et al., 2013). The associations between SA and GCA emerge during early postnatal development and remain throughout life (Walhovd et al., 2016). Very low birth weight children have smaller SA and lower GCA compared to those with normal birth weight (Solsnes et al., 2015). Another twin study showed that normative birth weight differences in monozygotic twins were related to differences in SA and GCA in adolescence (Raznahan et al., 2012).

In conclusion, the literature supports the importance of distinguishing between the genetically, ontogenetically, and phylogenetically distinct cortical metrics of SA and CT. Our results indicate that the well-replicated height-GCA association is accounted for by individual differences in cortical SA, and not CT, and that the genetic association between SA and GCA underlies the phenotypic height-GCA relationship.

## Acknowledgements

Supported by NIA R01 AG022381, AG018386, AG018384, AG050595 and R03 AG 046413.

